# High school Internship Program in Integrated Mathematical Oncology (HIP IMO) – five-year experience at Moffitt Cancer Center

**DOI:** 10.1101/2020.02.27.967950

**Authors:** Heiko Enderling, Philipp M. Altrock, Noemi Andor, David Basanta, Joel S. Brown, Robert A. Gatenby, Andriy Marusyk, Katarzyna A. Rejniak, Ariosto Silva, Alexander R.A. Anderson

**Affiliations:** Department of Integrated Mathematical Oncology, H. Lee Moffitt Cancer Center & Research Institute Tampa, FL, USA; Department of Cancer Physiology, H. Lee Moffitt Cancer Center & Research Institute Tampa, FL, USA; Department of Radiology, H. Lee Moffitt Cancer Center & Research Institute Tampa, FL, USA; University of South Florida, Morsani College of Medicine, Department of Oncologic Sciences, Tampa, Florida, USA

**Keywords:** Mathematical modeling, education, high school, cancer, oncology, interdisciplinary

## Abstract

Modern cancer research, and the wealth of data across multiple spatial and temporal scales, has created the need for researchers that are well-versed in the life sciences (cancer biology, developmental biology, immunology), medical sciences (oncology) and natural sciences (mathematics, physics, engineering, computer sciences). College undergraduate education is traditionally provided in disciplinary silos, which creates a steep learning curve at the graduate and postdoctoral levels that increasingly bridge multiple disciplines. Numerous colleges have begun to embrace interdisciplinary curricula, but students who double-major in mathematics (or other quantitative sciences) and biology (or medicine) remain scarce. We identified the need to educate junior and senior high school students about integrating mathematical and biological skills, through the lens of mathematical oncology, to better prepare students for future careers at the interdisciplinary interface. The High school Internship Program in Integrated Mathematical Oncology (HIP IMO) at Moffitt Cancer Center has so far trained 59 students between 2015 and 2019. We report here on the program structure, training deliverables, curriculum, and outcomes. We hope to promote such interdisciplinary educational activities early in a student’s career.

## Introduction

The H. Lee Moffitt Cancer Center (MCC) is a domestic, non-profit organization in Tampa, FL, USA. It is a free-standing institution and an NCI-designated comprehensive cancer center. The Center has over 7000 employees serving a population of over 21 million people. In 2008, MCC through the guidance of Robert Gatenby formed the Department of Integrated Mathematical Oncology, IMO. The central goal of the IMO is the integration of theoretical and computational modeling tools into clinical and experimental cancer research to aid in both the core understanding of cancer processes and the mechanisms that drive them [1]. By using a range of mathematical modeling approaches targeted at answering specific questions for different cancers the IMO aids in the elucidation of aspects of basic cancer biology, involving complex, dynamics systems (such as interactions between multiple cell types and micro-environment in formation of bone metastases) as well as development and testing of treatment strategies. This multi-model, multi-scale approach allows for a diverse and rich interdisciplinary environment that has created many novel approaches for the treatment and understanding cancer [2–23]. Crucial to our success is the true integration of theoretical models with cancer growth, evolution and treatment data, which requires sophisticated imaging techniques and potentially new experimental and clinical protocols. To this end the IMO is an interdisciplinary team of scientists incorporating experts in the field of mathematics, computer science, physics, genomics, evolutionary biology, ecology, imaging and visualization to name but a few. As of 2020, the IMO has eight faculty members, 14 postdoctoral fellows and six graduate students.

The *H*igh school *I*nternship *P*rogram in *I*ntegrated *M*athematical *O*ncology, HIP IMO, delivers interdisciplinary team science research experiences for high school students aged 16 or older by the time of the internship (http://www.moffitt.org/HIPIMO). This mentored summer training program is designed for motivated aspiring scientists to help prepare them for interdisciplinary cancer research careers. Working under the direction and guidance of faculty/scientist mentors in Moffitt’s IMO department, interns are involved in activities designed to foster the development of life-long research skills. Students are assigned individual research projects appropriate to their interests and abilities. The program runs for 8 weeks during the Hillsborough County public schools summer break. Applications are invited beginning November 1 for the following summer and close February 1, with enrollment decisions communicated by end of February of the internship year. Following a generous donation in 2017, HIP IMO students in the past two years received a $1,000 scholarship, additionally accommodation and travel support are provided for out of town students.

Students collaborate with an assigned mentor to create a research project with achievable goals in the time allotted, gain familiarity with standard methodologies in a safe environment, participate in scheduled lab meetings, acquire necessary data through experimentation, computation, surveys or other means, document findings in an appropriate format (laboratory journal, audiovisual recording or digital databases), review and discuss findings with research mentors, draw conclusions and make new plans, gain oral presentation experience by designing and delivering a 15-minute presentation to department members, Moffitt leadership, parents, teachers and patients at the HIP IMO research day, and write a three-page scientific report at the end of the program.

### Advertisement, application and student selection

The program is advertised as one of the research training programs at MCC on the MCC website. Additionally, in collaboration with Hillsborough County Public Schools administration, program announcements are distributed to local schools. While a large proportion of students learn about the program by word of mouth from friends and relatives of past participants, the program is widely advertised on social media (Twitter and LinkedIn). Applications can be downloaded in word and pdf format from the HIP IMO website, and include student information, voluntary reporting of demographics as well as school contact details. Applications do not request transcripts; rather each applicant has to answer three questions:

1. *Which, if any, college level biology, mathematics, or computer programming classes have you completed?*
2. *What research skills and expertise can you apply to the program? Please emphasize any cancer biology, mathematics, or computer programming experience that you may have*.
3. *What are your career interests/goals and how may the HIP-IMO program help your career?*

Applications include a letter of recommendation by at least one teacher. Students are selected by participating faculty members, and selection depends on the required skills and expertise for study the mentor wants to perform, and which student’s career goals best align with the project and focus of the research groups.

### Curriculum

After selection in the spring, the assigned faculty mentors communicate with their mentees on relevant literature and pre-requisites for successful completion of the summer internship. Each student is required to create a LinkedIn account and to join the HIP IMO group on LinkedIn to help tracking student careers after HIP IMO. The first week of HIP IMO is comprised of various boot camps to prepare the students for the hands-on modeling and simulation tasks of their projects. In the *cancer biology* and *cancer immunology* boot camps, students learn the molecular basis of cancer and immunology, the hallmarks of cancer [24,25] and the various cancer treatment strategies, including radiotherapy, chemotherapy, targeted therapy, and immunotherapy. With the establishment of the Center of Excellence for Evolutionary Therapy at Moffitt, students also receive training in adaptive cancer therapy [26–29]. In the *mathematical modeling* in cancer boot camp students learn the seven steps to modeling a biological problem (formulate question, determine basic model ingredients, describe system qualitatively, describe system quantitatively, analyze equations, checks and balances, relate results back to question; [30]), how to develop a flow diagram and construct a mathematical model from verbal descriptions of biological problems, and how to analyze such models.

As part of the boot camp we discuss different growth laws in oncology (linear, exponential, logistic, Gompertz [31,32]), predator-prey models in ecology and cancer [10,33], carcinogenesis, and cancer therapy [34,35]. The students also learn about the rigorous modeling pipeline of model calibration, validation and prediction performance evaluation [36]. To facilitate the programming core of HIP IMO, Matlab and Hybrid Automata Library (HAL) [37] boot camps are provided. Additional classes include biostatistics, medical physics, and intellectual property and licensing. After the one-week of boot camps, students perform individual mentored research in their assigned research group. Students’ are provided with a MacBook pro laptop with required software to execute and document their research. To facilitate networking and community building, every Friday we host a student-mentor luncheon where lunch is provided for all program participants.

In week two, students participate in a journal club and present a peer-reviewed publication that is relevant to their research project (5-10 minutes). Often this includes a paper authored by the faculty mentor. Students receive limited mentoring for the journal club presentation, to assess a variety of presentation styles. After all presentations, students select their favorite presentation and identify for themselves styles to adapt for their future presentations. Mentors help evaluate the individual presentations in their research groups. After four weeks, students present their research question and preliminary results in a midterm project presentation (5-10 minutes each).

The final highlight of the program is the HIP IMO research day on the last day of the internship (Friday of week eight). Each intern gives an oral presentation of their research project, with length dependent on the number of enrolled students (in total we aim to keep the total presentations under 4 hours). Each student is allowed to invite four family members to the research day. The program administration invites the teachers that provided the recommendation letter, as well as the school principals. The presentations are attended by all members of the IMO department, as well as MCC leadership and researchers. As with all presentations at the IMO, we extend a warm welcome to patient advocates to share their perspective and comment on the impact of each research project. After all presentations, the program provides a reception and dinner for students, families and researchers to network and to reflect upon the program.

### Outcomes

Since 2015, HIP IMO has enrolled 59 students (average 12/year) with an average enrollment rate of 31% (**Table 1**). 28 male (47%) and 31 female (53%) HIP IMO interns have graduated from the program, including six diversity students (10%). The majority of students (83%) came from Hillsborough County and other counties in Florida. Ten students (17%) came from out of state, including California (3), Ohio (2), Minnesota (1), New Jersey (1), Pennsylvania (1), North Carolina (1), Virginia (1). All students (20) who have graduated from high school since participation in HIP IMO continued their education in college, including Duke University (2), Columbia University (2), Rutgers University (2), University of Florida (UF, 3), University of South Florida (USF, 3), Georgia Tech (1), University of Missouri (1), University of Michigan (1), New York University (1), Vanderbilt University (1), Stanford University (1), UC Berkeley (1), UC Santa Barbara (1), The Ohio State University (1), Harvey Mudd College (1), Saint Louis University (1), Johns Hopkins University (1), Northwestern University (1), University of Miami (1). Seven students (35%) pursued a double major in the quantitative and life sciences, with computer sciences being the dominant major (6; 30%). Two students (10%) are pursuing a medical degree, and two students have graduated and continued in graduate school at the time of this report (Computer Science, Engineering). Two students (10%) have co-founded companies while in college. Two students have graduated from college and work as a software engineer.

**Table 1.**
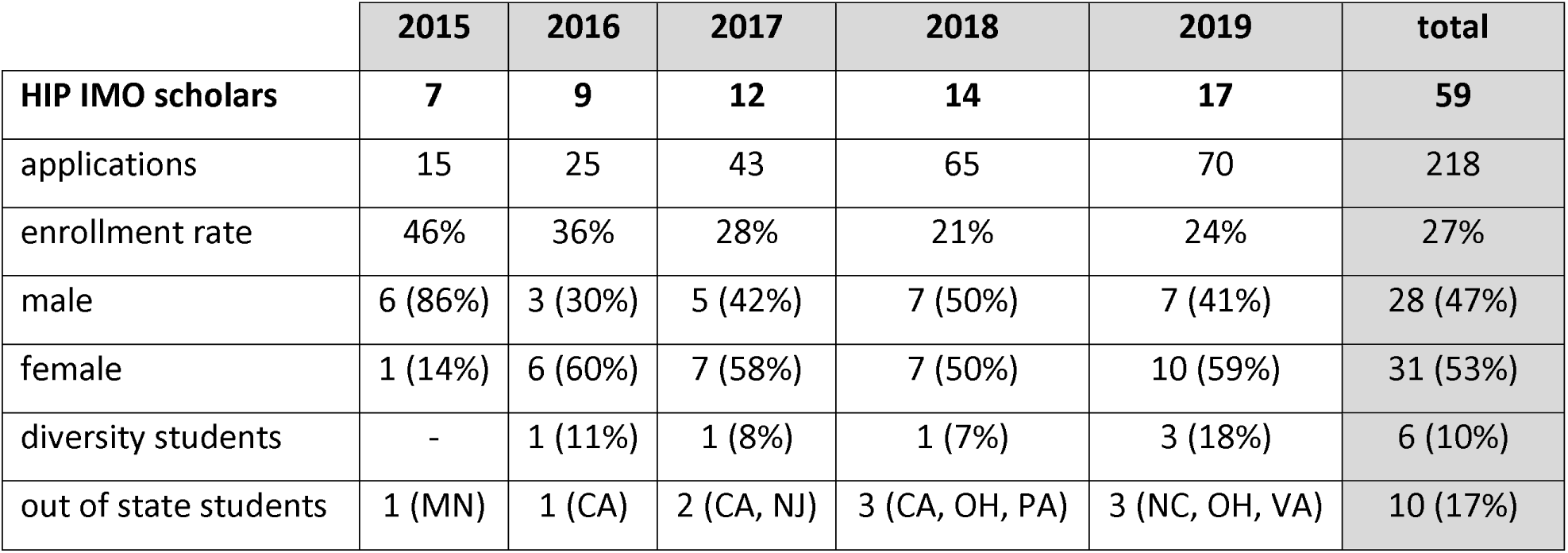
HIP IMO demographics 2015-2019.

|  | 2015 | 2016 | 2017 | 2018 | 2019 | total |
| --- | --- | --- | --- | --- | --- | --- |
| **HIP IMO scholars** | **7** | **9** | **12** | **14** | **17** | **59** |
| applications | 15 | 25 | 43 | 65 | 70 | 218 |
| enrollment rate | 46% | 36% | 28% | 21% | 24% | 27% |
| male | 6 (86%) | 3 (30%) | 5 (42%) | 7 (50%) | 7 (41%) | 28 (47%) |
| female | 1 (14%) | 6 (60%) | 7 (58%) | 7 (50%) | 10 (59%) | 31 (53%) |
| diversity students | - | 1 (11%) | 1 (8%) | 1 (7%) | 3 (18%) | 6 (10%) |
| out of state students | 1 (MN) | 1 (CA) | 2 (CA, NJ) | 3 (CA, OH, PA) | 3 (NC, OH, VA) | 10 (17%) |

HIP IMO has a defined a number of learning objectives that are assessed by self-evaluation in entry and exit surveys. Students rate their skills on a scale from one to ten (where one is the lowest and ten is the highest) in computer programming, mathematics, cancer biology, mathematical oncology, scientific research, literature review, oral presentation, and interdisciplinary team science. The average self-assessed skill level across all questions at the beginning of the program was 4.9 +/- 1.7 (n=59). By the end of the HIP IMO, the average response increased by 22% (6.0 +/- 1.8; n=49). The largest skill improvements were recorded in mathematical oncology (4.3 to 7.2, 69% increase, p<0.001 [Mann– Whitney U test]), computer programming (4.1 to 6.2, 52%, p<0.001), cancer biology (5.1 to 7.2, 42%, p<0.001) and mathematics (5 to 6.9, 38%, p<0.001; **Figure 1**). After HIP IMO, students strongly agreed *(i)* to have a network of peers and mentors that will help advance their careers (4.4 / 5) and *(ii)* to pursue a career in science (4.4 / 5). The majority of students identified to be interested in an interdisciplinary career in cancer research or medicine, which is reflected in the proportion of double majors students enrolled to in college (35%). When asked to describe HIP IMO with three adjectives the most common responses where fun (10), challenging (9), innovative (7), enlightening (7) and engaging (5) (**Figure 2**).

**Figure 1.**
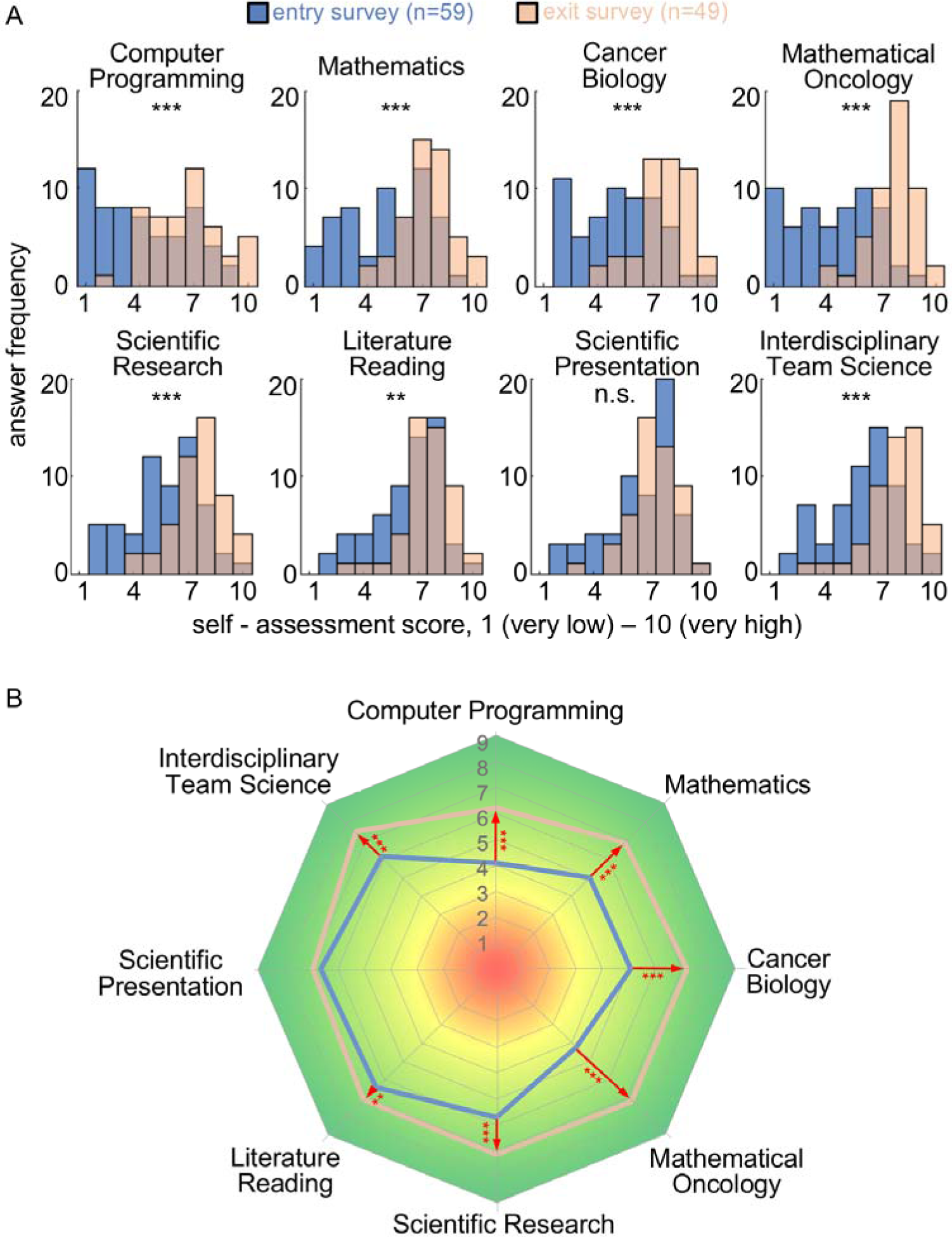
Self-reported skill assessment of students before (blue, n=59) and after HIP IMO (orange, n=49). **A**. Distributions of answers to surveys of self-assessment of different skills. Answers between 1- 10 (very low - very high). Statistical analysis of difference in response distributions using Mann-Whitney U test; ***indicate p<0.001; ** indicate p<0.01; n.s. not significant. **B**. Radar plot of entry and exit survey average responses to the self-evaluated skills demonstrating meeting of HIP IMO learning objectives.

**Figure 2.**
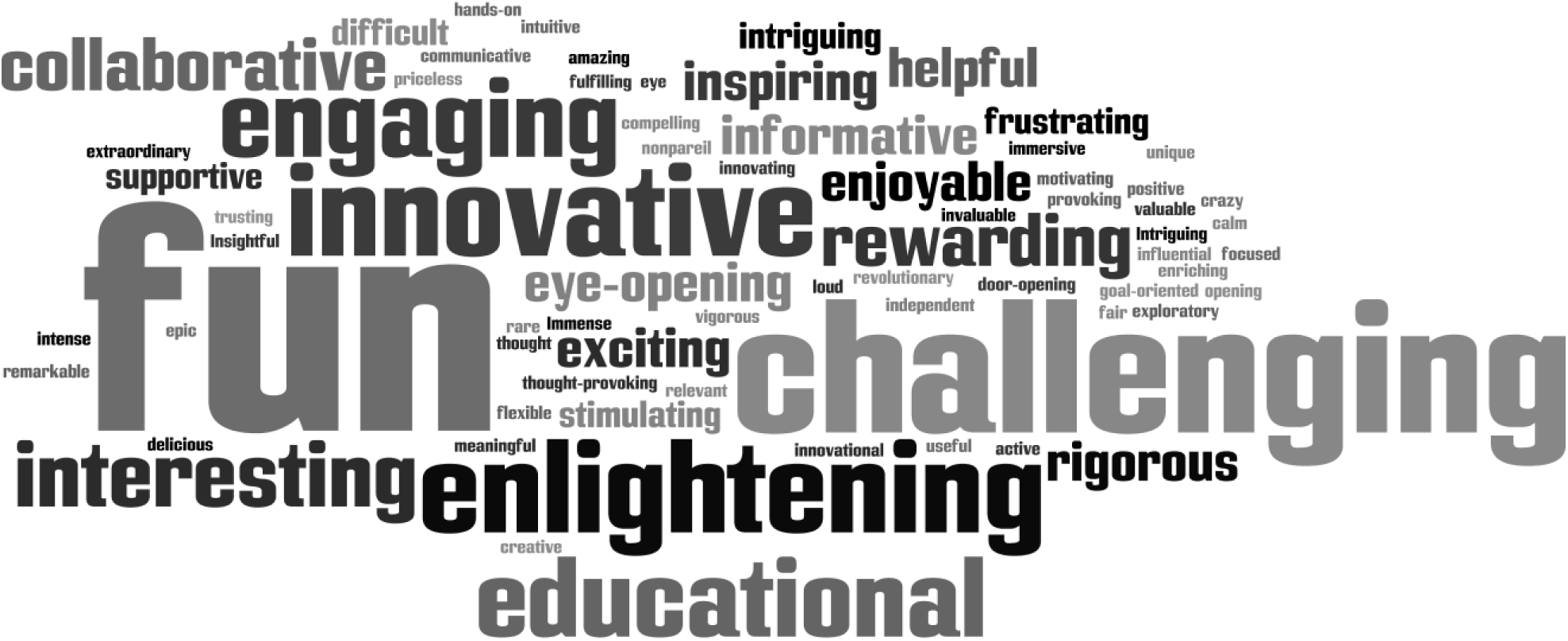
Word cloud of three adjectives students used to describe the program (N=49). Font size represents answer frequency.

Student projects contributed to a number of peer-reviewed and published articles and book chapters in topical and interdisciplinary journals including PLoS Computational Biology [38], Scientific Reports [39], International Journal of Radiation Biology [40], Games [41], Annals of Epidemiology [42], and Encyclopedia of Biomedical Engineering [43], as well as several BioRxiv preprints [44–47].

## Discussion and Conclusion

The first five years’ experience of the High school Internship Program in Integrated Mathematical Oncology at Moffitt Cancer Center has shown that junior and senior high school students of diverse demographic and socioeconomic backgrounds can perform excellent hands-on research in mathematical oncology. As demonstrated by the outcome analysis, learning objectives are clearly met under the guidance of dedicated faculty mentors and postdoctoral fellows.

The increasing number of interested students and outstanding applicants emphasize the need to educate high school students about interdisciplinary team science at the interface of mathematics and biology. The opportunity to explore career opportunities in the life sciences, and cancer research in particular, for students with a strong background and interest in mathematics, physics or computer science offers new interdisciplinary career trajectories for students before choosing college degrees in traditional discipline silos. Due to the strong commitment of faculty mentors, students can deliver quality research that may contribute to publications and grant proposals – clearly demonstrating a benefit to the teaching faculty as well. We hope that programs such as HIP IMO serve as motivation and blueprint for the development of similar programs to enable more high school students. These programs would enable a diverse group of students to get involved with cutting edge mathematical biology research that can directly influence and promote their future academic careers.

## Acknowledgements

HIP IMO is supported by NIH/NCI U54CA143970-05 (Physical Science Oncology Network) “Cancer as a complex adaptive system.” HIP IMO scholars are generously supported by the Richard O. Jacobson Foundation. In kind support is provided by the West Wing Hotel and the Museum of Sciences and Industry (MOSI). HIP IMO is grateful to Mehdi Damaghi, Asmaa El-Kenawi, Prabakaran Soundararajan, Dung-Tsa Chao, Michal Tomaszewski, Mark Robertson-Tesssi for teaching HIP IMO boot camps and classes, and the dedicated student and postdoctoral mentors for HIP IMO scholars (Etienne Baratchart, Renee Brady, Rafael Bravo, Ibrahim Chamsedine, Andrew Dhawan, Bina Desai, Meghan Ferrall-Fairbanks, Jill Gallaher, Daniel Glazar, Chandler Gatenbee, Rachel Howard, Aleksandra Karolak, Eunjung Kim, Gregory Kimmel, Anna Miller, Rafael Renatino Canevarolo, Jacob Scott,Praneeth Sudalagunta, Enakshi Sunassee, Robert Vander Velde, Jeffrey West, Mohammad Zahid). The authors are thankful to Daniel Glazar and Stefano Pasetto for statistical analysis of HIP IMO survey results. HIP IMO is especially thankful to Danae Paris for administrating the program, and support from the HCPS superintendent Jeff Eakins, HCPS Director for K-12 STEM Education Larry R. Plank, as well as Moffitt PSOC patient advocates Jeri Francoeur and Robert Butler. HIP IMO could not fully function without the critical infrastructure support from MCC and its continued commitment to educating the next generation of scientists.

